# Dorsomedial Striatal Glutamatergic Transmission Inhibits Binge Drinking in Selectively Bred Crossed High Alcohol Preferring Mice

**DOI:** 10.1101/2024.03.15.585024

**Authors:** Meredith R. Bauer, Megan M. McVey, Yanping Zhang, Stephen L. Boehm

**Affiliations:** Indiana Alcohol Research Center and Department of Psychology, Indiana University Indianapolis, Indianapolis, IN, 46202, USA

**Keywords:** Alcohol, AMPA Receptor, binge, cHAP, dorsomedial striatum

## Abstract

Crossed high alcohol preferring (cHAP) mice have been selectively bred to consume considerable amounts of alcohol resulting in binge drinking. The dorsal striatum (DS) is a brain region involved in action selection where the dorsomedial striatum (DMS) is involved in goal-directed action selection and dorsolateral striatum (DLS) is involved in habitual action selection. Alcohol use disorder (AUD) may involve a disruption in the balance between the DMS and DLS. While the DLS is involved in binge drinking, the reliance on the DMS and DLS in binge drinking has not been investigated in cHAP mice. We have previously demonstrated that glutamatergic activity in the DLS is necessary for binge-like alcohol drinking in C57BL/6J mice, another high drinking mouse. Because of this, we hypothesized that DLS glutamatergic activity would gate binge-like alcohol drinking in cHAP mice. cHAP mice underwent bilateral cannulation into the DMS or DLS and were allowed free-access to 20% alcohol for two-hours each day for 11 days. Mice were microinjected with the AMPA receptor (AMPAR) antagonist, NBQX, into the DMS or DLS immediately prior to alcohol access. AMPAR protein expression was also assessed in a separate group of animals in DS subregions following an 11-day drinking history. We found that intra-DMS (but not intra-DLS) NBQX, alters binge alcohol drinking, with intra-DMS NBQX increasing alcohol consumption. We also found that the ratio of GluA1 to GluA2 differs across DS subregions. Together, these findings suggest that glutamatergic activity in the DMS may serve to limit binge drinking in cHAP mice.

## Introduction

Alcohol related injury is one of the leading causes of preventable deaths in the United States (Pilar et al., 2020) and alcohol use can lead to the development of alcohol use disorder (AUD), a chronic, relapsing, hereditary disease (American Psychiatric Association, 2013). Additionally, frequency of binge drinking positively predicts the development of AUD, suggesting that binge drinking and AUD may be related (Addolorato et al., 2018). AUD may be associated with alterations in the balance in behavioral flexibility controlled by the dorsomedial and dorsolateral striatum (DLS). Further, the DLS is necessary for driving binge alcohol drinking in C57BL/6J mice (Bauer, McVey, & Boehm, 2022; Bauer et al., 2022; Haggerty et al., 2022). However, it is unknown whether selectively bred crossed high alcohol preferring mice also rely on the DLS for binge drinking or whether the balance in behavioral control between dorsal striatal subregions has shifted in the mouse model of a family history of AUD. Thus, the purpose of this project was to assess the involvement of the dorsomedial striatum (DMS) and the DLS in binge-like alcohol drinking using cHAP mice.

Binge drinking is characterized by an excessive high consumption of alcohol in a relatively short time window. It is defined by consuming 4 or more standard drinks in 2 hours for women or 5 or more drinks in two hours for males. Binge drinking can also be defined by reaching a blood alcohol concentration of above 80 mg/dl in just 2 hours. While many different genetic mouse models consume alcohol to the level of a binge, cHAP mice have been selectively bred to consume high amounts of alcohol (Oberlin et al., 2011; Matson et al., 2014). This selective breeding affords an opportunity to assess whether mechanistic changes in brain function occur as a function of a family history of high alcohol consumption which may offer insight into the neurobiology of AUD. For example, cHAP mice display compulsive-like alcohol drinking much faster and to more concentrated levels of quinine-adulterated alcohol (Houck et al., 2019; Sneddon et al., 2022), than other mouse models (see Table 1 in Bauer et al., 2021 for comparison). However, the neurobiological alterations that are the result a family history of high alcohol intake have not been fully characterized.

The dorsal striatum is a brain region most noted for its role in action selection where the DMS is important for goal-directed action selection and the DLS is important for habitual action selection for natural rewards (Yin et al., 2004; 2005) and for alcohol (Corbit et al., 2012; 2014). However, outside of action selection, the DMS and DLS are also important for alcohol drinking. For example, NMDA receptor antagonism in the DMS reduces alcohol drinking and an alcohol drinking history increases DMS AMPA receptor expression of GluA1 and GluA2 on the post-synaptic density within the DMS (Wang et al., 2012). Meanwhile, intra-DLS AMPA receptor antagonism reduces binge-like alcohol drinking (Bauer et al., 2022a, 2022b) in animals with a relatively short binge drinking history, but not long (i.e., three weeks of daily exposure). These data would suggest that both the DMS and DLS are necessary for alcohol drinking. However, none of these experiments assessed these behaviors in selectively bred high alcohol consuming lines. Filling this gap, Rangel-Barajas et al., 2021 recorded medium spiny neurons in the DMS and DLS of cHAP mice following chronic alcohol drinking. The authors found that an alcohol drinking history increases female sEPSC amplitude in the DMS and frequency in the DLS relative to water drinking controls suggesting alcohol drinking alters glutamatergic activity in both brain regions. Both the DMS and DLS are important for alcohol drinking and cHAP mice have altered glutamatergic activity in the DMS and DLS following an alcohol drinking history. However, it is not known whether selection for high alcohol consumption altered the function of the DMS and DLS for binge drinking. The purpose of this project was to determine if AMPA receptor antagonism of the DMS or DLS reduces binge-like alcohol drinking and whether an alcohol drinking history alters AMPA receptor expression in cHAP mice. Alterations in AMPA receptor expression may have functional implications for the glutamatergic synapse.

## Materials and methods

### Animals

Naïve adult male and adult female cHAP mice (n = 9/10 per sex) were bred on site at the IU Indianapolis School of Science, Indianapolis, IN (for more information on the selection and characterization of cHAP mice, please see Oberlin et al., 2011). Animals were individually housed in a vivarium with 12-hour:12-hour reverse light–dark cycle for at least 1 week prior to the start of experiments. Mice were housed in nonfiltered wired top standard shoe box mouse cages (18.4 cm wide, 29.2 cm long, 12.7 cm tall), and were given food (Lab Diet 5001, Rodent Diet) and tap water ad libitum with the exception of water bottle removal during the DID sessions. Procedures were approved by the IUI School of Science Institutional Animal Care and Use Committee and conformed to the Guide for the Care and Use of Laboratory Animals (The National Academic Press, 2003).

### Bilateral Cannulation Surgery

The DMS and DLS were targeted for bilateral cannulations using a Kopf stereotaxic alignment system. All surgical procedures have been described in depth elsewhere (Bauer et al., 2022a, Bauer et al., 2022b). Briefly, DMS coordinates used were M/L: +/-1.25 mm, A/P: + 0.5 mm, and D/V: - 2.75 mm and DLS coordinates were M/L: +/-2.5 mm, A/P: + 0.38 mm, and D/V: - 3.0 mm. Following surgery, mice received a single subcutaneous injection of 5 mg/mL carprofen (a post-operative pain treatment) at 10 mL/kg and were placed a heating pad until recovery from anesthesia (approximately 30-minutes). Mice were given ad-libitum food and water and at least a week of recovery prior to DID. Following the completion of the behavioral data collection, brains were taken for verification of cannula placement. Mice were humanely euthanized, brains were extracted, and flash-frozen in 2-methylbutane at -20 to -40 degrees C. Brain slices were stained with cresyl violet and placements were determined using the Paxinos and Franklin brain atlas (2001) as a reference.

### Stylet Changing and Microinjections

Stylets were changed daily. Mice were given mock injections on day 5 immediately prior to DID by using microinjectors that extended 0.5 mm past the cannula and restraining the mouse for the duration of a microinjection (2.5 minutes). Mice were microinjected on days 7, 9, and 11 with one of three drug concentrations (saline (control), 0.15 μg/side, or 0.5 μg/side NBQX) into the DMS or DLS immediately prior to DID in a within-subjects, Latin-square design. On days 6, 8, and 10 mice were not microinjected but DID access to alcohol was still provided. Drug concentrations were chosen based on previous findings in our group (Bauer et al., 2022a; 2022b) and others (Corbit et al., 2012; 2014; & Wang et al, 2012)

### Drinking-in-the-Dark (DID)

DID is a limited-access model of binge-like alcohol intake which has been described in-depth elsewhere (Theile, Crabbe, Boehm, 2014). Briefly, mice get access to a ball-bearing sipper tube filled with 20% v/v alcohol (190 proof alcohol purchased from Pharmco, Inc (Brookfield, CT) mixed with tap water). This is administered in their home cage for two-hours, three-hours into the dark cycle each day. cHAP mice, though selectively bred for high alcohol intakes in a two-bottle choice drinking procedure, readily binge drink under the DID paradigm (Ardinger et al., 2020; Winkler & Grahame, 2023). DID occurred for a total of 11 days. Consumption was measured by reading the sipper tubes to the nearest 0.025 mL. Volumes were adjusted for leak based on the volume leaked from a tube in an empty cage on the same rack.

### Blood Alcohol Concentrations (BACs)

Retro-orbital sinus bloods were collected immediately following DID on the final infusion day. BACs were acquired using an Analox EtOH Analyzer (Analox Instruments, Lunenburg, MA as previously described (Bauer et al., 2021). Some (∼1/2) of the bloods from the DMS experiment are missing due to equipment malfunction.

### Locomotor Activity

Locomotor activity was monitored in the home cage during DID sessions on infusion days. Locomotor activity was collected using the Opto M-3 system (Columbus Instruments) which collects locomotor activity by summating beam breaks over a set time-interval from infrared beams that surround the perimeter of the home cage, described in depth here (Bauer, Garcy, & Boehm, 2020).

### Subcellular Fractionation of Tissue

A separate group of mice were given an 11 day DID history of either water or alcohol (20%). 24-hours after DID on day 11, brains were harvested, and tissue was dissected from the DMS and DLS. We then followed a membrane fractionation protocol adapted from Dunah & Standaert, 2001. Tissue was removed from freezer and transferred to wet ice where we added 400 μL ice-cold TEVP-320 buffer (320mM sucrose). Tissue homogenization is done with TissueTearor for 10-second pulses at up to 50% power for 3 cycles with 5 second intervals. Centrifuge homogenate for 10 min at 800 x g, 4°C to separate supernatant and pellet. Transfer supernatants to a new tube and centrifuge at 9,500 x g, 4°C for 15 minutes. Wash pellet once in TEVP-35.6 (35.6mM sucrose) and centrifuge briefly, and discard the washing buffer. Resuspend pellets in 75uL TEVP+ with protease & phosphatase inhibitors by brief vortex. The protein was quantified on 1:2 dilutions. These samples were then used for western blot.

### Western Blot

Using the subcellular fractionations described above, protein concentrations were determined using the Bio-Rad Protein Assay Kit (Bio-Rad protein assay Dye reagent concentrates). Samples were counterbalanced by sex and drinking history across gels for Western-Blot. Briefly, 40 microgram of sample protein was 95°C heat denatured for 5 min in a final volume of 30 ul containing 4x Loading Dye and 5% of 1M DTT. The protein was separated by gel electrophoresis using 4%-20% gradient gel (Bio-Rad) running at 120V in the 1X Tris/Glycine/SDS buffer (Bio-Rad). The gel is transferred using Trans-Blot turbo midi nitrocellulose trasfer membrane (Bio-Rad) for 7 minutes at 2.5A/25V. Blocking the membrane was done with 5% nonfat milk in TBS Buffer for 1 hr. The primary antibody was dilution 1:1000 with 5% nonfat milk in 1x TBS with 0.1% Tween 20, and incubation at 4°C overnight. (The Anti-Glutamate Receptor 1 (AMPA subtype), Rabbit polyclonal Antibody, cat # ab31232, from abcam. The Anti Glur2 ployclonal antibody, cat# PA5-19496 form Thermo fisher Scintific). The membrane was washed with TBS-T (TBS+ 0.1% Tween 20) for 3 times, 10 min each. Adding secondary antibody at 1:10000 dilutions and incubated for 1 hour at room temperature.v(IRDye 800 CW Coat anti-Rabbit IgG (H+L) from LI-COR. cat# 926-32211). The membrane was Washed 3 times, 10 min each. Document and analyze the blot using CLx Odyssey scanner and software Image Studio Lite Ver 5.2. The results was normalized with beta-actin

### Experimental Design and Statistical Analyses

Statistical analyses (RM ANOVA, Pearson’s correlation, or t-tests) were conducted for alcohol drinking, locomotor activity, BACs, and molecular assays. For all analyses, sex was used as a factor. When male and female mice did not differ from one another in any analyses, we collapsed across sex. Greenhouse-Geisser corrections were applied as necessary. Differences were considered significant at p < 0.05. Figures were made using GraphPad Prism 8 and BioRender (BioRender.com). Data were analyzed using R Studio (www.r-project.org).

## Results

The effect of intra-DLS NBQX is shown in Figure 1. A timeline for this part of the experiment can be seen above Figure 1. Baseline alcohol drinking (days 1–5) for male and female cHAP mice with DLS cannula are shown in Figure 1B (placements shown in Figure 1A). Alcohol consumption is displayed in grams consumed per kilogram of body weight. RM one-way ANOVA of baseline alcohol drinking across day revealed no significant differences of day (p > 0.05). On days 7, 9, and 11 mice were microinjected with one of three concentrations of NBQX. RM one-way ANOVA revealed that intra-DLS NBQX did not alter alcohol consumption (p > 0.05), Figure 1C. Immediately following DID on day 11, blood samples were collected. Pearson’s correlation indicated that BACs positively and significantly predicted alcohol intake, r(21) = 0.78, p < 0.0001, Figure 1D. One-way ANOVA assessing BAC for each concentration of NBQX revealed no significant differences (p > 0.05). BAC mean ± SEM for saline = 88.18 ± 33.79, 0.15 μg/side = 106 ± 30.81, 0.5 μg/side = 96.19 ± 31.25 (data not shown). Locomotor activity was assessed during the DID session immediately following infusions. RM two-way ANOVA of time and drug concentration revealed a main effect of time, F(5.7, 125) = 5.2, p < 0.0001) and no other significant differences or interactions (p > 0.05), Figure 1E. Average locomotor activity from the entire DID session following infusions was analyzed. RM one-way ANOVA revealed no significant differences (p > 0.05).

**Figure 1.**
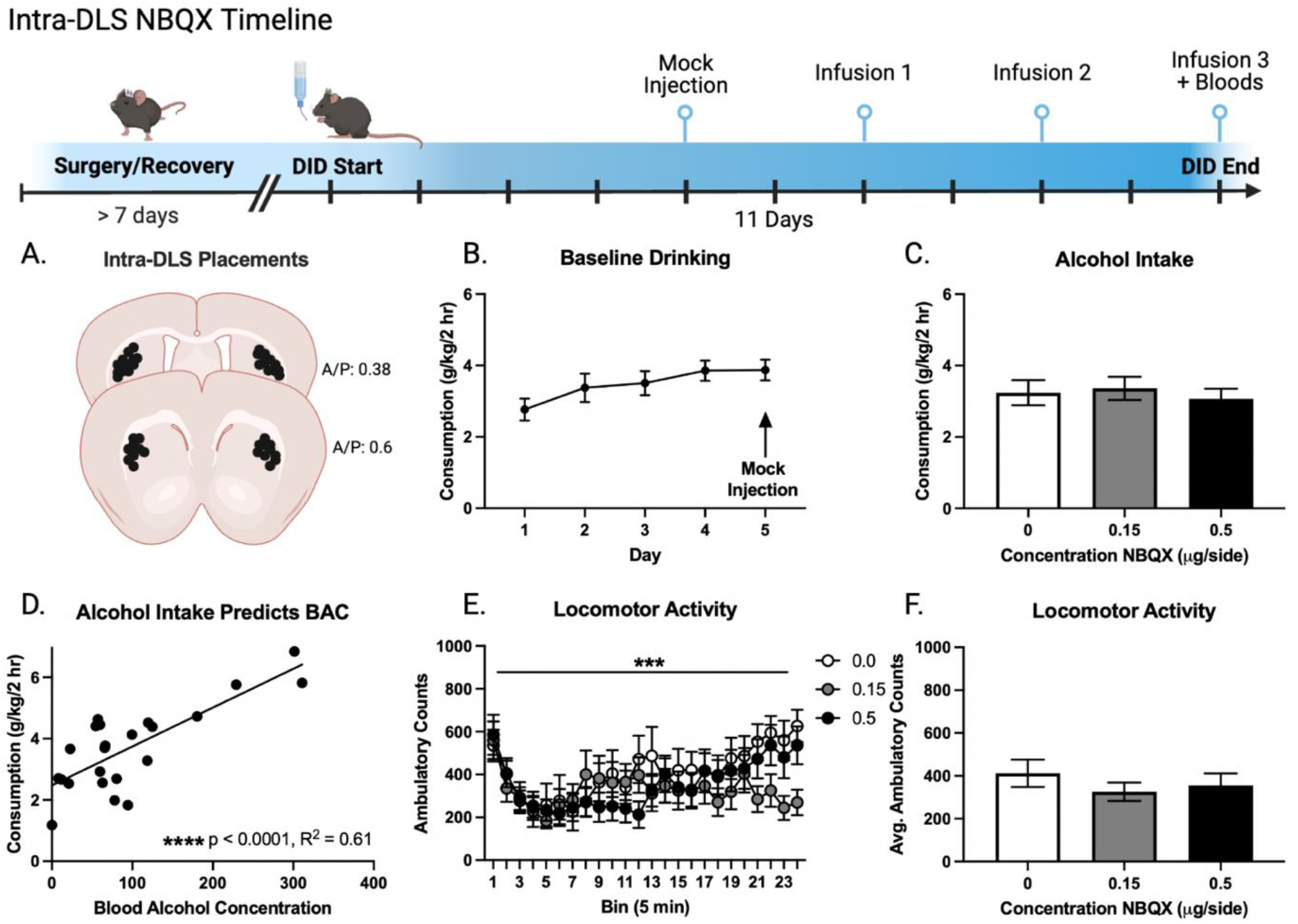
Effect of Intra-DLS NBQX on Alcohol Drinking. All mice underwent bilateral cannulation surgery, recovered, and began DID. Microinjections of NBQX were given immediately prior to DID. Retro-orbital sinus bloods were collected immediately following DID on day 11. **A**. Representative placements of DLS cannulation. Placements were compared with Paxinos and Watson’s Mouse Brain Atlas. **B**. Average daily alcohol intake. RM one-way ANOVA revealed no effect of day (p > 0.05). **C**. Effect of intra-DLS NBQX on alcohol consumption. RM one-way ANOVA revealed no effect of drug concentration (p > 0.05). **D**. Pearson’s correlation revealed that alcohol intake positively predicted BACs (****p < 0.0001). **E**. Locomotor activity during DID sessions following infusions. RM two-way ANOVA revealed a main effect of time (***p < 0.001). **F**. Total average locomotor activity during the DID sessions following infusions. RM one-way ANOVA revealed no effect of drug concentration (p > 0.05). Data are displayed as mean ± standard error of the mean.

The effect of intra-DMS NBQX is shown in Figure 2. A timeline for this part of the experiment can be seen above Figure 1. Baseline alcohol drinking (days 1–5) for male and female cHAP mice with DMS cannula are shown in Figure 2B (placements shown in Figure 2A). Alcohol consumption is displayed in grams consumed per kilogram of body weight. RM one-way ANOVA of baseline alcohol drinking across day revealed no significant differences of day (p > 0.05). On days 7, 9, and 11 mice were microinjected with one of three concentrations of NBQX. RM one-way ANOVA revealed that intra-DMS NBQX significantly increased alcohol consumption, F(2,32) = 4.77, p = 0.015, η^2^_G_ = 0.072, Figure 2C. Post-hoc Bonferroni corrected paired t-test revealed that 0.15 μg/side NBQX significantly increased intake compared with saline (p = 0.047). Immediately following DID on day 11, blood samples were collected. 8 samples were not analyzed from this group due to equipment malfunction. Pearson’s correlation indicated that BACs positively and significantly predicted alcohol intake, r(10) = 0.90, p < 0.0001, Figure 2D. We were not able to assess BACs across each drug concentration due to low power due to missing several blood samples. Locomotor activity was assessed during the DID session immediately following infusions. RM two-way ANOVA of time and drug concentration revealed a main effect of time, F(6.5, 116.8) = 2.6, p = 0.02 and no other significant differences or interactions (p’s > 0.05), Figure 2E. Average locomotor activity from the entire DID session following infusions was analyzed. RM one-way ANOVA revealed no significant differences (p > 0.05), Figure 2F.

**Figure 2.**
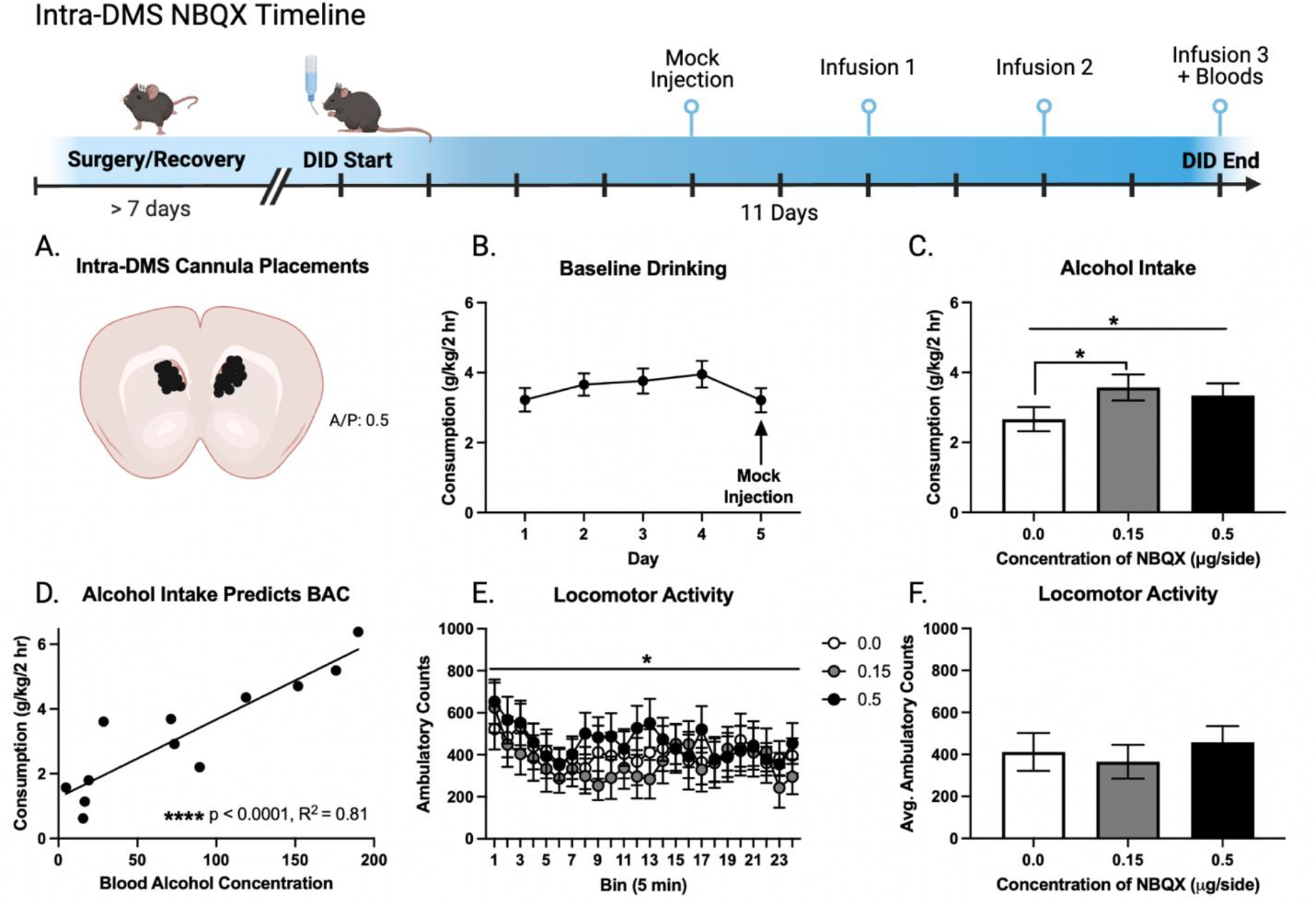
Effect of Intra-DMS NBQX on Alcohol Drinking. All mice underwent bilateral cannulation surgery, recovered, and began DID. Microinjections of NBQX were given immediately prior to DID. Retro-orbital sinus bloods were collected immediately following DID on day 11. **A**. Representative placements of DMS cannulation. Placements were compared with Paxinos and Watson’s Mouse Brain Atlas. **B**. Average daily alcohol intake. RM one-way ANOVA revealed no effect of day (p > 0.05). **C**. Effect of intra-DMS NBQX on alcohol consumption. RM one-way ANOVA revealed a main effect of drug concentration (p > 0.05). Post-hoc Dunnet’s corrected paired t-test revealed a significant difference between 0 and 0.15 μg/side. **D**. Pearson’s correlation revealed that alcohol intake positively predicted BACs (****p < 0.0001). **E**. Locomotor activity during DID sessions following infusions. RM two-way ANOVA revealed a main effect of time (*p < 0.05). **F**. Total average locomotor activity during the DID sessions following infusions. RM one-way ANOVA revealed no effect of drug concentration (p > 0.05). Data are displayed as mean ± standard error of the mean.

The effect of alcohol or water drinking on post-synaptic density (PSD) GluA1 and GluA2 expression is shown in Figure 3. A timeline for this part of the experiment is shown above Figure 3. Baseline alcohol drinking (days 1–11) for male and female cHAP mice is shown in Figure 3A. Alcohol consumption is displayed in grams consumed per kilogram of body weight. RM two-way ANOVA of baseline alcohol drinking across day in male and female cHAP mice revealed a main effect of sex, F(1,14) = 49.9, p < 0.001, η^2^_G_= 0.29 and a main effect of day, F(10,140) = 4.10, p= 0.001, η^2^_G_= 0.21, Figure 3A. Baseline water drinking (days 1-11) for male and female cHAP mice is shown in Figure 3B. RM two-way ANOVA of baseline water drinking across day in male and female cHAP mice revealed no significant effects (p’s > 0.05). Images of gels for the DMS and DLS protein expression for male and female cHAP mice is shown in Figure 3C. The effect of alcohol or water drinking on DLS PSD GluA1 and GluA2 expression is shown in Figure 3D. RM three-way ANOVA of drinking history, sex, and protein revealed a significant main effect of protein F(1,54) = 33.60, p < 0.001, η^2^_G_ = 0.38, Figure 3D, with the expression of GluA2 being higher than that of GluA1. The effect of alcohol or water drinking on DMS PSD GluA1 and GluA2 expression is shown in Figure 3E. RM three-way ANOVA of drinking history, sex, and protein revealed no significant effects (p’s > 0.05), Figure 3E. The effect of GluA1/A2 ratio on brain region (DLS vs DMS) is shown in Figure 3F. Unpaired t-test comparing GluA1/A2 ratio for the DLS and DMS revealed a significantly higher ratio for the DMS relative to the (****p < 0.0001), Figure 3F.

**Figure 3.**
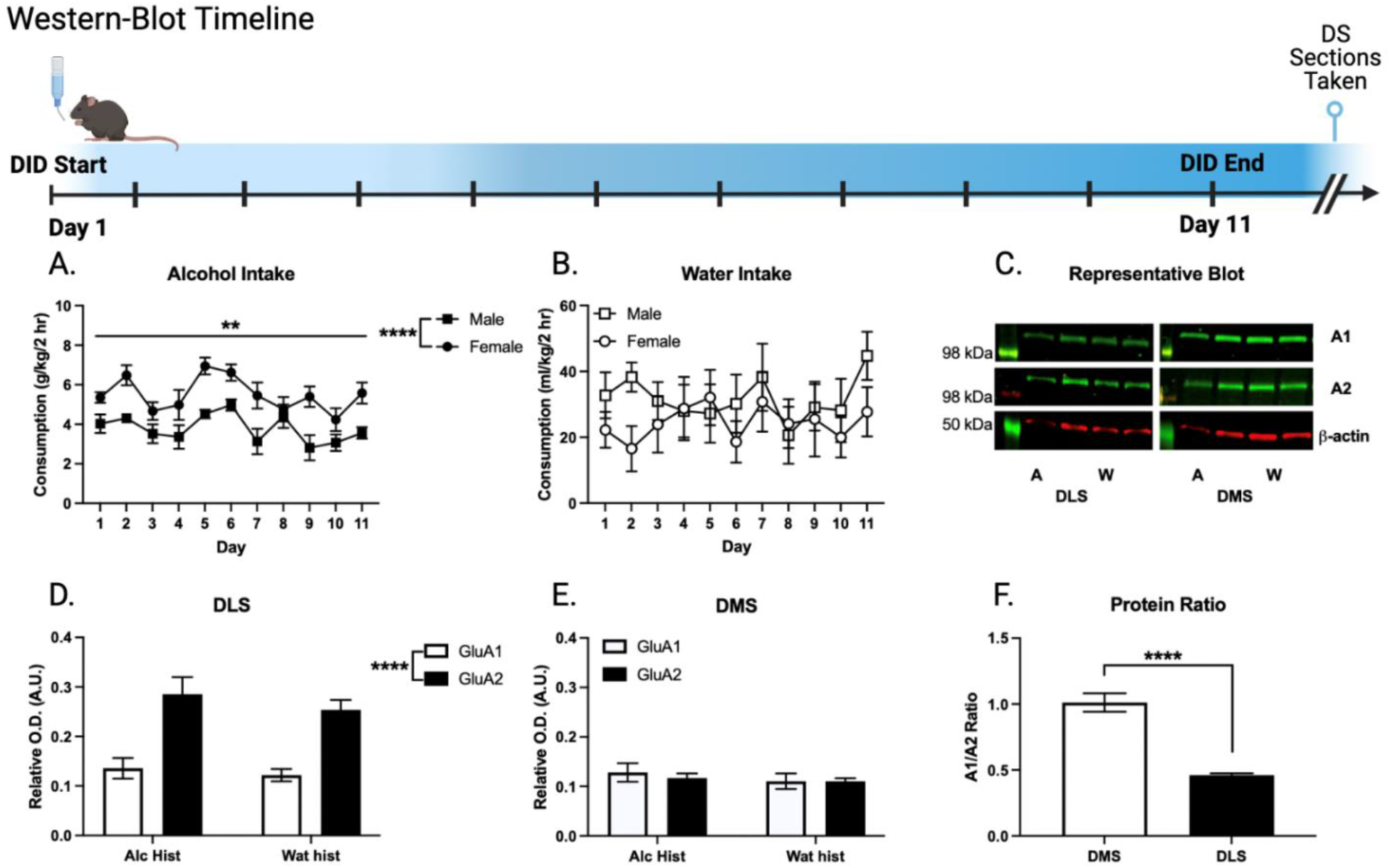
Dorsal Striatum PSD-Enriched GluA1 and GluA2 Protein Expression. All mice underwent DID of either alcohol or water. 24-hours later DS sections were harvested for western-blot. **A**. Baseline alcohol drinking in male and female mice. RM two-way ANOVA revealed a main effect of sex (****p < 0.0001) and day (**p < 0.01). **B**. Baseline water drinking in male and female control mice. **C**. Representative blots. The far left of each image is the ladder for each gel with molecular weight of the protein. Treatment groups from left to right: male alcohol, female alcohol, male water, female water from the DLS and male alcohol, female alcohol, male water, female water from the DMS. These are shown for GluA1 (top), GluA2 (middle) which were normalized to beta-actin (bottom). **D & E**. GluA1 and GluA2 protein expression from DLS or DMS tissue in alcohol and water history mice. Two-way ANOVA revealed a main effect of protein (****p < 0.0001) in DLS, but not DMS tissue. **F**. Ratio of GluA1/GluA2 in DLS vs DMS. Unpaired t-test revealed a significant difference (****p < 0.0001). Data are displayed as mean ± standard error of the mean.

## Discussion

The goal of this project was to probe the relationship between glutamatergic activity in the dorsal striatum and selection for high alcohol preference in male and female cHAP mice. Based on previous research, we hypothesized that intra-DMS or -DLS AMPAR antagonism would reduce binge like alcohol drinking. Counter to our hypothesis we found that intra-DMS AMPAR receptor antagonism increased alcohol drinking while intra-DLS AMPAR antagonism did not alter binge alcohol drinking. Further, we found that the DMS has a significantly higher ratio of GluA1/GluA2 on PSD enriched fractions than the DLS, regardless of alcohol drinking history. We used male and female cHAP mice but did not find any significant interactions or main effects of sex as it relates to glutamatergic activity. However, female cHAP mice drank significantly more than male cHAP mice in the western-blot experiment. These data provide a novel finding that manipulation of glutamatergic activity in the DMS causes binge-like alcohol drinking and contribute to the growing literature on sex differences in alcohol related behaviors.

The finding that non-specific AMPAR antagonism in the DMS increased alcohol drinking is to our knowledge, the first of its kind. This is not to say that manipulations in the DMS have never increased drinking, but that this sort of global inhibition has not shown such an effect before. For example, NMDA receptor antagonism, which is a similar approach to our global inhibition, decreased alcohol drinking (Wang et al., 2010), though intake was dependent on operant alcohol deliveries which were also reduced by NMDA receptor antagonism. However, studies of cell or input specific targets in the DMS show similar results to our DMS finding. DMS D1 spiny neuron activation or stimulation of cortical terminals into the DMS increases alcohol consumption (Cheng et al., 2017; Ehinger et al., 2021; & Ma et al., 2021) though controls for nonspecific ligand/light activation were not controlled for so interpretation is limited. In line with the previously described findings, Lrrk2 (a D1 receptor negative modulator) knock out mice which show increased D1 receptor expression in the dorsal striatum drink more alcohol and more quinine-adulterated alcohol (a measure of compulsive-like alcohol drinking) than wild-type controls. These data suggest that our non-specific inhibition of DMS AMPA receptors causing an increase in drinking may be working to potential D1 neurons. Though the current manuscript offers no direct evidence of this, it is possible that inhibiting DMS AMPARs caused an increase in D1 activity or a decrease in D2 activity which could in turn cause increased D1 activity, ultimately resulting in an increase in alcohol drinking. Future research should investigate the cellular mechanism at work.

While we found no differences in PSD enriched GluA1 or GluA2 across drinking history (alcohol or water history) we did find that the DMS has a significantly higher GluA1/GluA2 ratio compared with the DLS. One important consideration with this finding is that the ratio is higher on the PSD which is functionally important for plasticity. This could suggest that the DMS innately expresses more GluA1 on the PSD than the DLS which may be related to specific signaling metrics (e.g., greater reliance on calcium permeable AMPAR expression). For example, AMPA receptors are most commonly tetrameric dimers of GluA1/A2 pairs (Henley, 2016) and are sodium permeable. However, GluA2 lacking AMPARs become calcium permeable. The most common form of CP-AMPARs are GluA1 homomers (Henley, 2016). Thus, it is possible that the DMS of cHAP mice depend more on CP-AMPAR mechanisms to prevent drinking. Thus, our pharmacological blockade of AMPARs may have targeted an abundance of CP-AMPARs working to prevent drinking thus causing a release of a brake, resulting in an increase in alcohol drinking. It is likely that cell type is important for interpretation of this ratio difference. The DMS is composed of D1 and D2 medium spiny neurons and several interneuron types (GABAergic and muscarinic). Previous research has found that D1 agonists, D2 antagonists, D1 and D2 coactivation, and muscarinic acetylcholine antagonism each increase striatal S845 phosphorylation (Xue et al., 2017) which is important for synaptic insertion of the GluA1 subunit (Man, Sekine-Aizawa& Huganir, 2007). Thus, knowledge of which cell type(s) are experiencing increased GluA1 expression is likely important for further interpretation of this finding.

We found that intra-DLS AMPAR antagonism did not alter alcohol drinking. This finding is counter to our central hypothesis which was based on previous research from our lab and others. We have twice replicated that intra-DLS AMPAR antagonism reduces binge alcohol drinking after a short, but not extended drinking history in C57BL/6J mice (Bauer et al., 2022a; 2022b). Specifically, after C57BL/6J mice drank for three weeks, which we have previously shown results in compulsive quinine-resistant alcohol drinking, they were no longer sensitive to intra-DLS AMPAR antagonism (Bauer et al., 2021). Since cHAP mice demonstrate innate compulsive alcohol drinking (Houck et al., 2019) and the dorsal striatum is highly relevant for compulsive alcohol use (Everitt & Robbins, 2016), is it possible that selection for high alcohol drinking in cHAP mice resulted in alterations in reliance on the dorsal striatal relative to C57BL/6J mice for alcohol drinking behavior (i.e., binge and compulsive alcohol drinking).

Finally, we adequately powered our experiments to be able to detect sex differences, yet we did not see many sex specific effects. Female mice (C57BL/6J, high alcohol preferring (HAPs), or cHAP) typically drink more alcohol than their male counterparts (Bauer, Garcy, Boehm, 2020; Bauer, McVey, & Boehm, 2021; Winkler & Grahame, 2023), though not always (Ardinger et al., 2021; Sneddon et al., 2022; Bauer et al., 2022a; 2022b). We found, what we have seen before, that cannulated mice do not show sex differences in basal alcohol drinking (Figures 1B and 2B). However, female mice drank more than male mice in the western blot experiment (Figure 3A and 3B). We did not observe sex differences in microinjection or western-blot results suggesting no sex effects in AMPA-specific manipulations. These data contribute to the growing body of knowledge on sex differences in alcohol behaviors.

The findings in this project are limited in the following ways. First, we used a non-specific manipulation of glutamatergic activity which essentially makes our experiment a brain region specific pharmacological lesion study. Because of our choice of manipulation, we are unable to glean insight onto which cells or inputs are being affected to alter drinking. However, while our technical choice is limited, it also provides a unique opportunity to better understand the function of the DMS or DLS in binge-like alcohol drinking. We found that intra-DMS AMPAR antagonism increases binge-alcohol drinking and that intra-DLS AMPAR antagonism does not alter alcohol drinking in cHAP mice. We hypothesized that intra-DMS AMPAR or intra-DLS AMPAR antagonism would reduce binge-like alcohol drinking. However, much of our hypotheses were based on pharmacological lesioning studies assessing the DMS or DLS’ role in alcohol seeking or drinking in an operant setting. For example, Wang et al, 2012 found that intra-DMS NMDA receptor antagonist reduced responding for alcohol and alcohol drinking in an operant setting in which alcohol access is contingent on pressing a lever for access. Because of this, if seeking is reduced, drinking inherently has to be because reduced seeking reduces access to alcohol. In line with this, much of the literature dissecting the DMS or DLS focus on goal-directed or habitual responding for alcohol (Corbit, Nie, Janak et al., 2012; 2014; Fanelli et al., 2013; Renteria et al., 2018; 2022l; Ma et al., 2022; Baltz et al., 2023; Schreiner et al., 2023). Given the known differences in alcohol seeking versus taking behavior (Samson, Slawecki, Sharpe, & Chappell, 1998; Samson & Czachowski, 2003), experiments like the one detailed in our project are necessary to further understand the neurobiology of alcohol drinking. Our aim to identify the function of the DMS and DLS for binge drinking in cHAP mice adds to this literature.

Another limitation is we did not directly compare cHAPs with another high drinking mouse such as C57BL/6J. We are interpreting our findings in the context of previous literature using other high drinking mice but do not directly investigate the effect of selective breeding for high alcohol consumption. Previous research comparing high alcohol preferring lines with other high alcohol consuming mice within experiment have found both similarities and discrepancies between genetic mouse models. For example, systemic administration of NBQX, the same AMPAR antagonist used in this project, reduced binge drinking in C57BL/6J mice but not in selectively bred high alcohol preferring (HAP) mice (Bauer, Garcy, & Boehm, 2020), systemic baclofen reduced binge drinking in both C57BL/6J and HAP mice, but in different ways (Bauer, Hernández, Kasten, & Boehm, 2022), and McCane et al., 2018 found that systemic administration of a catechol-O-methyl transferase inhibitor reduced alcohol drinking in selectively bred alcohol preferring rats (P-rats) but not Wistar rats, even though both rat genotypes consume alcohol. This limitation should be considered when interpreting the findings in this paper.

In summary, the findings of this project provide novel insight into the function on the dorsal striatum for binge alcohol drinking in male and female cHAP mice. We demonstrate that whereas pharmacological blockade of DMS AMPARs increases binge alcohol drinking, pharmacological blockade of DLS AMPARs does not. We also demonstrate that the GluA1/A2 ratio is greater in the DMS than the DLS of cHAP mice. These findings differ from what has previously been observed in C57BL/6J mice which may be the result of selective breeding for high alcohol consumption. More broadly, we find that DMS glutamatergic activity acts to reduce alcohol drinking in cHAP mice and that GluA1/A2 ratios differ between the DMS and DLS. Future research should investigate the cell type specific mechanisms at work.

## Notes

This work was supported by the Indiana Alcohol Research Center and NIH/NIAAA grants AA07611, T32AA07462, and 1F31AA029273-01A1.

The authors report no conflicts of interest.

### Competing Interest Statement

The authors have declared no competing interest.

## References

Addolorato, G., Vassallo, G. A., Antonelli, G., Antonelli, M., Tarli, C., Mirijello, A., … & Gasbarrini, A. (2018). Binge drinking among adolescents is related to the development of alcohol use disorders: results from a cross-sectional study. Scientific reports, 8(1), 12624.

Ardinger, C. E., Winkler, G., Lapish, C. C., & Grahame, N. J. (2021). Effect of ketamine on binge drinking patterns in crossed high alcohol-preferring (cHAP) mice. Alcohol (Fayetteville, N.Y.), 97, 31–39. 10.1016/j.alcohol.2021.09.004

Baltz, E. T., Renteria, R., & Gremel, C. M. (2023). Chronic alcohol exposure differentially alters calcium activity of striatal cell populations during actions. Addiction Neuroscience, 100128.

Bauer, M. R., Hernández, M., Kasten, C. R., & Boehm II, S. L. (2022). Systemic administration of racemic baclofen reduces both acquisition and maintenance of alcohol consumption in male and female mice. Alcohol, 103, 25–35.

Bauer, M. R., McVey, M. M., & Boehm, S. L. (2021). Three weeks of binge alcohol drinking generates increased alcohol front‐loading and robust compulsive‐like alcohol drinking in male and female C57BL/6J mice. Alcoholism: Clinical and Experimental Research, 45(3), 650–660.

Bauer MR, McVey MM, Boehm SL 2nd. Drinking History Dependent Functionality of the Dorsolateral Striatum on Gating Alcohol and Quinine-Adulterated Alcohol Front-Loading and Binge Drinking [published online ahead of print, 2022 Oct 11]. Alcohol. 2022;S0741-8329(22)00099-4. doi:10.1016/j.alcohol.2022.09.005

Bauer, M. R., McVey, M. M., Germano, D. M., Zhang, Y., & Boehm II, S. L. (2022). Intra-dorsolateral striatal AMPA receptor antagonism reduces binge-like alcohol drinking in male and female C57BL/6J mice. Behavioural brain research, 418, 113631.

Bauer MR, Garcy DP, Boehm SL 2nd. Systemic Administration of the AMPA Receptor Antagonist, NBQX, Reduces Alcohol Drinking in Male C57BL/6J, But Not Female C57BL/6J or High-Alcohol-Preferring, Mice. Alcohol Clin Exp Res. 2020;44(11):2316–2325. doi:10.1111/acer.14461

Corbit LH, Nie H, Janak PH. Habitual alcohol seeking: time course and the contribution of subregions of the dorsal striatum. Biol Psychiatry. 2012;72(5):389–395. doi:10.1016/j.biopsych.2012.02.024

Corbit LH, Nie H, Janak PH. Habitual responding for alcohol depends upon both AMPA and D2 receptor signaling in the dorsolateral striatum. Front Behav Neurosci. 2014;8:301. Published 2014 Sep 2. doi:10.3389/fnbeh.2014.00301

Dunah AW, Standaert DG. Dopamine D1 receptor-dependent trafficking of striatal NMDA glutamate receptors to the postsynaptic membrane. J Neurosci. 2001;21(15):5546–5558. doi:10.1523/JNEUROSCI.21-15-05546.2001

Everitt, B. J., & Robbins, T. W. (2016). Drug addiction: updating actions to habits to compulsions ten years on. Annual review of psychology, 67, 23–50.

Fanelli, R. R., Klein, J. T., Reese, R. M., & Robinson, D. L. (2013). Dorsomedial and dorsolateral striatum exhibit distinct phasic neuronal activity during alcohol self‐ administration in rats. European Journal of Neuroscience, 38(4), 2637–2648.

Oberlin, B., Best, C., Matson, L., Henderson, A., & Grahame, N. (2011). Derivation and characterization of replicate high-and low-alcohol preferring lines of mice and a high-drinking crossed HAP line. Behavior genetics, 41, 288–302.

Haggerty, D. L., Munoz, B., Pennington, T., Di Prisco, G. V., Grecco, G. G., & Atwood, B. K. (2022). The role of anterior insular cortex inputs to dorsolateral striatum in binge alcohol drinking. Elife, 11, e77411.

Henley, J. M., & Wilkinson, K. A. (2016). Synaptic AMPA receptor composition in development, plasticity and disease. Nature Reviews Neuroscience, 17(6), 337–350

Houck, C. A., Carron, C. R., Millie, L. A., & Grahame, N. J. (2019). Innate and acquired quinine‐ resistant alcohol, but not saccharin, drinking in crossed high–alcohol‐preferring mice. Alcoholism: Clinical and Experimental Research, 43(11), 2421–2430.

McCane, A. M., DeLory, M. J., Timm, M. M., Janetsian-Fritz, S. S., Lapish, C. C., & Czachowski, C. L. (2018). Differential COMT expression and behavioral effects of COMT inhibition in male and female Wistar and alcohol preferring rats. Alcohol, 67, 15–22.

Ma, T., Huang, Z., Xie, X., Cheng, Y., Zhuang, X., Childs, M. J., … & Wang, J. (2022). Chronic alcohol drinking persistently suppresses thalamostriatal excitation of cholinergic neurons to impair cognitive flexibility. The Journal of Clinical Investigation, 132(4)

Matson, L. M., Kasten, C. R., Boehm, S. L., & Grahame, N. J. (2014). Selectively bred crossed high‐alcohol‐preferring mice drink to intoxication and develop functional tolerance, but not locomotor sensitization during free‐choice ethanol access. Alcoholism: Clinical and Experimental Research, 38(1), 267–274.

Man, H. Y., Sekine-Aizawa, Y., & Huganir, R. L. (2007). Regulation of α-amino-3-hydroxy-5-methyl-4-isoxazolepropionic acid receptor trafficking through PKA phosphorylation of the Glu receptor 1 subunit. Proceedings of the National Academy of Sciences, 104(9), 3579–3584.

Paxinos G, Franklin KBJ. The Mouse Brain in Stereotaxic Coordinates, 2nd ed., Academic Press, San Diego, 2001.

Renteria, R., Baltz, E. T., & Gremel, C. M. (2018). Chronic alcohol exposure disrupts top-down control over basal ganglia action selection to produce habits. Nature communications, 9(1), 211.

Renteria, R., Cazares, C., Baltz, E. T., Schreiner, D. C., Yalcinbas, E. A., Steinkellner, T., … & Gremel, C. M. (2021). Mechanism for differential recruitment of orbitostriatal transmission during actions and outcomes following chronic alcohol exposure. Elife, 10, e67065.

Samson, H. H., Slawecki, C. J., Sharpe, A. L., & Chappell, A. (1998). Appetitive and consummatory behaviors in the control of ethanol consumption: a measure of ethanol seeking behavior. Alcoholism: Clinical and Experimental Research, 22(8), 1783–1787.

Samson, H. H., & Czachowski, C. L. (2003). Behavioral measures of alcohol self-administration and intake control: rodent models.

Schreiner, D. C., Wright, A., Baltz, E. T., Wang, T., Cazares, C., & Gremel, C. M. (2023). Chronic alcohol exposure alters action control via hyperactive premotor corticostriatal activity. Cell Reports, 42(7).

Sneddon, E. A., Schuh, K. M., Fennell, K. A., Grahame, N. J., & Radke, A. K. (2022). Crossed high alcohol preferring mice exhibit aversion-resistant responding for alcohol with quinine but not footshock punishment. Alcohol, 105, 35–42.

Thiele TE, Crabbe JC, Boehm SL 2nd. “Drinking in the Dark” (DID): a simple mouse model of binge-like alcohol intake. Curr Protoc Neurosci. 2014;68:9.49.1-9.49.12. Published 2014 Jul 1. doi:10.1002/0471142301.ns0949s68

Wang, J., Hamida, S. B., Darcq, E., Zhu, W., Gibb, S. L., Lanfranco, M. F., … & Ron, D. (2012). Ethanol-mediated facilitation of AMPA receptor function in the dorsomedial striatum: implications for alcohol drinking behavior. Journal of Neuroscience, 32(43), 15124–15132.

Winkler, G. A., & Grahame, N. J. (2023). Home Cage Voluntary Alcohol Consumption Increases Binge Drinking without Affecting Abstinence-Related Depressive-Like Behaviors or Operant Responding in Crossed High Alcohol-Preferring (cHAPs). Alcohol.

Xue, B., Chen, E. C., He, N., Jin, D. Z., Mao, L. M., & Wang, J. Q. (2017). Integrated regulation of AMPA glutamate receptor phosphorylation in the striatum by dopamine and acetylcholine. Neuropharmacology, 112(Pt A), 57–65. 10.1016/j.neuropharm.2016.04.005

Yin, H. H., Knowlton, B. J., & Balleine, B. W. (2004). Lesions of dorsolateral striatum preserve outcome expectancy but disrupt habit formation in instrumental learning. European journal of neuroscience, 19(1), 181–189.

Yin, H. H., Knowlton, B. J., & Balleine, B. W. (2005). Blockade of NMDA receptors in the dorsomedial striatum prevents action-outcome learning in instrumental conditioning. The European journal of neuroscience, 22(2), 505–512. doi:10.1111/j.1460-9568.2005.04219.x

